# Inorganic profiles of preimplantation embryos and the association of zinc with *Nanog* expression in the blastocyst

**DOI:** 10.1101/2025.05.14.654061

**Authors:** Julia L. Balough, Thomas V. O’Halloran, Francesca E. Duncan, Teresa K. Woodruff

## Abstract

Elements such as iron, copper and zinc play essential roles in the mammalian oocyte, egg, and embryo, however among these metals, zinc plays unique regulatory roles. Temporal fluctuations in zinc concentrations drive reproductive milestones such as meiotic resumption, egg activation, and initiation of the mitotic cell cycle. Roles for zinc in late preimplantation embryo development, have not been well characterized. Using a quantitative element approach we report the inorganic profiles of mouse embryos progressing through the late blastocyst stage. We find that blastocysts, like oocytes and eggs, and distinct to somatic cells, maintain higher levels of zinc than copper and iron. All three of these essential metals are more abundant in the inner cell mass, which contains the population of pluripotent stem cells that give rise to the fetus, relative to the trophectoderm which gives rise to the placenta and extraembryonic tissues. To test whether zinc abundance was associated with mitotic progress and cell fate lineage, we perturbed zinc homeostasis during blastocyst formation by artificially raising intracellular zinc concentrations with zinc pyrithione. This treatment during the morula-to-blastocyst transition when cell fate lineages emerge resulted in an elevation of zinc in the ICM. This treatment did not impact cell number but did increase expression of the pluripotency and epiblast marker, *Nanog*. These results demonstrate that the inorganic profiles of the late preimplantation embryo retain elemental hallmarks of earlier developmental stages and perturbation of zinc levels alters pluripotency gene expression in the blastocyst.

**In Brief:** Zinc plays important regulatory roles during oocyte maturation, fertilization and early embryo development; however, the role of zinc in the late preimplantation embryo is unknown. In this study, inorganic profiling of mouse embryos reveal zinc levels are higher than copper and iron in the late blastocyst and zinc supplementation during the morula-to-blastocyst alters expression of pluripotency marker, *Nanog*.

## Introduction

Transition metals, such as Mn, Fe, Co, Cu, Zn and Mo, play highly specific structural and catalytic roles that are necessary for proper cellular functioning (Eide, 2006, Uriu-Adams and Keen, 2010). Among these metals, zinc plays a variety of unique physiological roles in control of developmental processes. Highly localized fluctuations in zinc content drive regulatory events during maturation and activation of the female gamete. For instance, spatial-temporal fluctuations in zinc concentrations in the oocyte regulate the switch between meiotic arrest and resumption in the fertilized egg and the initiation of mitosis in the preimplantation embryo (Kim et al., 2010, Bernhardt et al., 2011, Kim et al., 2011, Bernhardt et al., 2012, Kong et al., 2012, Tian et al., 2014, Kong et al., 2015, Duncan et al., 2016, Zigo et al., 2022, Chen et al., 2023). These reproductive cell types have unique inorganic signatures relative to somatic cell types in that zinc is the most abundant transition metal, with the total atom level existing at an order of magnitude higher than other elements such as iron and copper (Kim et al., 2010, Kong et al., 2015).

A suite of single cell analysis methods, including synchrotron-based x-ray microscopy (XFM), as well as small molecule perturbations of metal content have been essential to establishing these roles for zinc. For example, the concentration of zinc increases by 50% during meiotic maturation, and this rise is essential for the oocyte to reach metaphase of meiosis II (MII). Preventing this zinc accrual with a zinc chelator, N,N,N′,N′-tetrakis-(2-pyridylmethyl)-ethylenediamine (TPEN), results in an arrest at a telophase I-like stage (Kim et al., 2010, Kim et al., 2011). In contrast to the increase of zinc that occurs during meiotic maturation, fertilization triggers a decrease via the exocytosis of 10 billion zinc atoms in what has been termed the “zinc spark” (Kim et al., 2011, Duncan et al., 2016). If the fertilized egg fails to release zinc at this stage or if zinc content is experimentally elevated using a zinc ionophore, zinc pyrithione (ZPT), eggs do not complete meiosis, fail to progress into mitosis, or begin to reform the metaphase II spindle and actin cap (Kim et al., 2011). Following fertilization, zinc levels remain relatively stable, and higher than both copper and iron, throughout 8-cell stage of preimplantation embryo development (Kong et al., 2015). Chelation of zinc in cleavage stage embryos, specifically in 2- and 4-cell embryos, results in developmental arrest due to failed mitosis, altered chromatin structure, and inhibition of global transcription (Kong et al., 2015). These findings demonstrate an important function of zinc in early preimplantation embryo development, but its role in later stages, namely the morula and blastocyst stages, has not been systematically investigated. We hypothesize that if zinc dynamics are necessary for early embryonic development, then zinc levels may play an essential role in blastocyst formation in the late preimplantation embryo in the mouse.

There are multiple cellular events that occur during preimplantation embryo development, including the maternal-to-zygotic transition, compaction and polarization of internal and external cell populations, and cell lineage specification (Zernicka-Goetz et al., 2009, Jedrusik, 2015). The blastocyst is the result of the first cell fate decision in mammals. In early embryo development, the blastomeres can become all cell and tissue types; however, as the embryo develops through the cleavage stages and morula, this potential is lost and the blastomeres segregate into different fates (Suwinska et al., 2008, Maemura et al., 2021). The blastocyst contains two cell populations: the inner cell mass (ICM) which will give rise to all embryonic cell types and the trophectoderm (TE) cells which develop into extraembryonic cell types such as the placenta (Zernicka-Goetz et al., 2009). The ICM and TE cell populations are characterized by exclusive expression of several transcription factors. *Cdx2,* caudal type homeobox 2, is a gene specifically expressed in the TE that inhibits the pluripotency marker *Oct4*. *Oct4* (*Pou5f1)* is a POU-domain transcription factor highly expressed in the ICM. *Oct4* expression is necessary for induction of pluripotency and upregulates pluripotent maintenance factor, *Nanog,* while inhibiting expression of *Cdx2 (Zernicka-Goetz et al., 2009, Wu et al., 2010).* The second differentiation step in embryo development is specification within the ICM into the epiblast (EPI) and primitive endoderm (PE). The EPI maintains pluripotency through expression of *Nanog* and gives rise to all fetal tissue types whereas the PE expresses differentiation markers, *Gata6, Gata4* and *Sox17*, and gives rise to extra-embryonic cells populations such as the visceral and parietal endoderm which facilitate nutrient and metabolic embryo support prior to implantation. Development of ICM lineages is necessary as loss of either EPI or PE can result in developmental arrest and lethality (Artus and Chazaud, 2014). The mechanisms of cell partitioning and cell fate determination in the preimplantation embryo are essential for blastocyst formation, implantation, and fetal development.

In this study, we defined the elemental content of that late preimplantation embryo utilizing synchrotron-based XFM and assessed the effect of elevated zinc during blastocyst development. We demonstrated that total zinc levels are constant from the 8-cell through blastocyst stages, and that zinc content is significantly higher in each of these stages compared to iron and copper. Further we observed a striking enrichment of zinc atoms in the ICM of the blastocyst. To determine whether zinc homeostasis is important for blastocyst formation, we utilized zinc pyrithione (ZPT) treatment to elevate zinc content during the morula-to-blastocyst transition and evaluated its impact on blastocyst formation. We defined a minimal level of ZPT treatment that effectively increased zinc. While ZPT had no effect on total, ICM, or TE cell number in the resulting blastocysts, this treatment did induce an increase in pluripotency marker gene expression. These results support the importance of zinc homeostasis during late preimplantation embryo development and suggest a regulatory role in cell identity within the ICM.

## Materials and Methods

### Animals and breeding

This study was approved by Northwestern University’s Institutional Animal Care and Use Committee (IACUC). Female CD1 mice, aged 6-8 weeks, were obtained from Envigo (Madison, WI, USA) and housed in a controlled barrier facility at Northwestern University’s Center for Comparative Medicine (Chicago, IL, USA). Humidity, temperature and photoperiod (14 light: 10 dark) were kept constant and food (Teklad Global irradiated chow 2916) and water were provided ad libitum. To maximize embryo yield, female mice were hyperstimulated by intraperitoneal injection of 10 IU pregnant mare’s serum gonadotropin (PMSG, EMD Bioscience, San Diego, CA, USA). 44-46 hours later female mice were superovulated with 10 IU human chorionic gonadotropin (hCG, Sigma-Aldrich, St. Louis, MO, USA). Next, two female mice were paired with one male CD1 mouse (aged 10-25 weeks) for trio breeding. For all experiments described in this study, a biological replicate consisted of pooled embryos, derived from cohorts of 10 female mice, which were randomly distributed into the control or treatment groups. This process was repeated for each experimental replicate.

### Embryo collection and preparation for X-ray fluorescent microscopy (XFM)

Eight-cells, morulae, and blastocysts were collected from mated females at 72, 96 and 120 hours post-hCG, respectively. The oviduct and uteri of bred female mice were collected into dissection media containing Leibovitz’s L-15 medium (L-15, Sigma Aldrich, St. Louis, MO, USA) supplemented with 0.1% polyvinyl alcohol (PVA, Sigma Aldrich, St. Louis, MO, USA) and cut with scissors to release the embryos. Embryos were collected and washed in pre-equilibrated (37LJC, 5% CO_2_) potassium complex optimized media (KSOM, EMD Millipore, Burlington, MA, USA). Early blastocysts were characterized by a blastocoel cavity <50% of total embryo volume and late blastocysts had blastocoels >50% total embryo volume. Embryos were prepared for XFM as previously described (Kim et al., 2010, Kong et al., 2015). Briefly, embryos were washed three times in 100 mM, pH 7 ammonium acetate to remove metal salts. Next embryos were loaded on a 2 mm x 2 mm silicon nitride window (Norcada Inc., Canada), excess media was removed by wicking and embryos air dried and were stored at room temperature until imaging.

### X-ray fluorescence imaging and quantification

XFM was performed at the Advanced Photon Source on beamline 2-ID-E at the Argonne National Laboratory as previously described (Kong et al., 2015). In brief, 10 keV X-rays were monochromatized and focused to a spot size of 0.5 μm x 0.5 μm using Fresnel zone plate optics. Raster scans were done using fly-scan mode with a 0.5-μm step size and a dwell time of 50 ms. Fluorescence spectra were collected using a silicone drift detector (Vortex-ME4). Fluorescence spectra were integrated along the z-axis, calibrated using thin film standard from AXO (AXO DRESDEN, Dresden, Germany) and two-dimensional elemental maps were built for each element to display elemental concentration in units of μg/cm^2^. Image processing and analysis were performed using MAPS 1.8.0.00 software. The elemental content was determined for specific regions of interest (ROI) using the phosphorus channel to capture: whole embryo, ICM, TE, as well as background, which was subtracted after normalization for area (Chen et al., 2023).

### Zinc pyrithione treatment

Previous studies have shown egg cytotoxicity upon treatment with ZPT (Sigma-Aldrich, St. Louis, MO, USA) concentrations of 10, 20 or 50 µM sometimes with 5-10 minute treatment window (Taki et al., 2004, Kim et al., 2011, Bernhardt et al., 2012). To establish an optimal concentration of ZPT that increased intracellular zinc but did not induce toxicity, a 10-1000 nM dose response embryo culture assay was performed. A 50 mM stock solution of ZPT in dimethyl sulfoxide (DMSO, Sigma-Aldrich, St. Louis, MO, USA) was made and diluted as described below. Female mice were hyperstimulated, superovulated and mated as previously described. 36 hours after pairing for trio breeding, two-cell embryos were collected from the oviducts of mated females into Leibovitz’s L-15 medium (L-15, Sigma Aldrich, St. Louis, MO, USA) supplemented with 0.1% polyvinyl alcohol (PVA, Sigma Aldrich, St. Louis, MO, USA). Any unfertilized, fragmented or unsynchronized embryos were discarded. Two-cell embryos were cultured for 48 hours in Advanced KSOM (EMD Millipore, Billerica, MA, USA) in an incubator (37LJC, 5% CO_2_) until the morula stage. Because the treatment time was increased to 24 hours to capture morula to blastocyst transitions, the dose response assay was performed with ZPT at concentrations of 0 nM (vehicle control) (n= 8 trials), 10 nM (n=4 trials), 100 nM (n=7 trials) and 1 μM (n=5 trials). Morulae (n=10/dose) were cultured in KSOM with control or ZPT for 24 hours and assessed for blastocyst development. Live embryos were imaged on brightfield on EVOS FL Auto Cell Imaging System (Thermo Fisher Scientific, Waltham, MA) at 10X. The maximum concentration that showed no statistical difference in embryo development was 100 nM ZPT and was used for all subsequent experiments. The DMSO concentration was matched and underwent identical serial dilution from the ZPT stock (50 mM) to ensure control and treatment had the same DMSO concentration in the final solution. Therefore, there was 1:500,000 or 0.000002% (v/v) DMSO in the vehicle or treatment (100 nM) final media.

### Labile zinc labeling and live-cell imaging

To assess if ZPT treatment led to an increase in intracellular zinc content, labile zinc concentration was evaluated after treating live embryos with the vital zinc-responsive fluorescent probe ZincBY-1 (Que et al., 2015). Vehicle or ZPT treated blastocysts were incubated for 30 minutes prior to imaging with 50 nM ZincBY-1, a labile zinc-specific fluorophore, and 1 μg/ml Hoechst 33342 (Thermo Fisher Scientific). Blastocysts were rinsed and loaded into pre-equilibrated KSOM drops under mineral oil (Irvine Scientific, Santa Ana, CA, USA) on a glass bottom dish (MatTek Corporation, Ashland, MA, USA). Confocal microscopy was performed on a Leica TCS SP5 inverted laser-scanning confocal microscope (Leica Microsystems, Heidelberg, Germany) at 10X air objective. All exposure and microscope settings including laser power, gain, pinhole and zoom were set to the ZPT treatment group and kept constant for all embryos. Z-stacks were taken of each embryo with a step size of 2.39 μm. Image analysis for signal intensity was performed in ImageJ (NIH, Bethesda, MD, USA). Z-stack image files were compressed into a single maximum projection image. Using the threshold tool, maximum and minimum threshold for each embryo were selected and total intensity was measured. The absolute intensity of ZincBY-1 was determined by dividing the raw intensity by the embryo area (µm^2^).

### Quantitative real-time PCR

For qPCR of cell fate markers, 3-5 experimental replicates were performed for a total of 300-500 blastocysts (100 blastocysts per biological replicate with 50 in control and 50 in treatment group). For qPCR of zinc transporters, a single experimental replicate (100 blastocysts) was performed with 50 blastocysts in the control and 50 blastocysts from the treatment group. Blastocysts were derived from 10 female mice per replicate and randomly sorted into experimental groups. Within each qPCR biological replicate, three technical replicates were performed and averaged. After 24-hour culture with control of ZPT, blastocysts were isolated and lysed in 50 μl of extraction buffer from the Arcturus PicoPure RNA Isolation Kit (Thermo Fisher Scientific) and snap frozen and stored at -80°C until RNA isolation. RNA isolation was performed per manufacturers recommendations with on-column DNase I treatment. Eluted RNA was reverse transcribed, and cDNA was synthesized using SuperScript III First-Strand cDNA synthesis kit with random hexamers (Thermo Fisher Scientific). Real time PCR was performed using SYBR green master mix and primers (Supplemental Table 1) (Integrated DNA Technologies) on the StepOnePlus real time PCR system (Applied Biosystem) with 56°C annealing temperature and 40 run cycles. Each PCR plate run included 3 internal replicates per treatment and gene. Changes in expression were expressed as fold change using the 2^−ΔΔCt^ method and normalized to *B-actin* expression. To ensure β-actin was suitable to use as a reference gene, we calculated the standard deviation, the coefficient of variance, and the maximum fold change of CT values for β-actin across all experimental replicates and found a SD of 0.17, CV of 0.73% and MFC of 1.01 indicating β-actin expression was stable in our samples (Pfaffl et al., 2004, Willems et al., 2006, de Jonge et al., 2007, Mamo et al., 2007).

### Blastocyst immunocytochemistry

Blastocysts were fixed in 3.8% paraformaldehyde for 1 hour then washed through blocking buffer (PBS containing 0.3% BSA, 0.01% Tween-20 and 0.02% NaN_3)._ Blastocysts were permeabilized in 1X PBS, 0.3% BSA, 0.1% Triton X-100, and 0.02% NaN_3_ for 20 minutes then washed through blocking buffer. Embryos were incubated overnight at 4LJC in mouse monoclonal anti-CDX2 (1:100, BioGenex, MU392A) and rhodamine phalloidin (1:200, Thermo Fisher Scientific, R415). Embryos were washed through blocking buffer and incubated in Alexa Fluor-488 donkey anti-mouse IgG (1:200, Invitrogen, A21202) secondary antibody for 1 hour at 37LJC. Embryos were washed through blocking buffer and moved through a Vectashield+DAPI gradient (25%, 50%, 75% in blocking buffer) then mounted in 100% Vectashield+DAPI (H-1200, Vector Laboratories, Inc.). Images were taken using a Leica TCS SP5 inverted laser-scanning confocal microscope (Leica Microsystems) at 40X oil immersion objective. Z-stacks were taken at 2.39 μm section thickness. Images were processed using ImageJ (NIH, Bethesda, MD, USA). For cell counting, newly appearing nuclei were annotated on each stack of the total z-stack. Total embryo cells were determined by counting DAPI or nuclei, TE cells were CDX2+ nuclei and ICM were CDX2-nuclei. Mitotic cells were determined by counting cells in metaphase or anaphase per total cells counted.

### RNAscope

RNAscope was performed according to manufactures instructions using the RNAscope Multiplex Fluorescent v2 Assay and protocol as previously described (Xie et al., 2018). In brief, blastocysts were fixed in 100 μl of 3.8% paraformaldehyde and 0.1% polyvinylpyrrolidone overnight. To detect TE and ICM transcripts, probes synthesized against the mouse *Cdx2* (438921), *Pou5f1(Oct4)* (426961), *Nanog* (501891) and *Gata4* (417881) were used (Advanced Cell Diagnostics). All reagents were provided with the RNAscope Multiplex Fluorescent v2 Assay. Opal fluorophores 520. 570, 690 (Akoya Biosciences, Marlborough, MA) were used as secondary antibodies. Embryos were imaged on the Leica TCS SP5 inverted laser-scanning confocal microscope (Leica Microsystems) at 40X oil immersion objective. Z-stacks were taken at 2.39 μm section thickness. Transcripts per embryos were captured by measuring intensity using ImageJ (NIH, Bethesda, MD, USA). The absolute intensity for each transcript was determined by dividing the raw intensity by the embryo area (µm^2^). To capture the number of transcript abundant nuclei per embryo, z-stacks each embryo were max projected. The area of *Nanog* or *Gata4* enriched transcript was traced and the number of nuclei marked by DAPI within that region was counted and normalized to the area of the whole embryo (µm^2^).

### Statistical analysis

Statistical analysis was performed using Graphpad Prism 9 software. Data was tested for normal distribution with the Shapiro-Wilk test. When appropriate multiple comparisons were tested with one-way ANOVA or Kruskal-Wallis ANOVA. For a single comparison, Mann-Whitney test was used. Values are expressed in mean ± standard deviation unless otherwise stated. A P-value <0.05 was considered significant.

## Results

### Iron, copper and zinc content across the late preimplantation embryo

To determine the elemental composition of zinc, iron, and copper throughout late preimplantation embryo development, we performed synchrotron-based XFM on 8-cells, morulae, early and late blastocysts derived from *in vivo* matings (Fig. 1A, Supp. Figs. 1-4). Like the oocyte, egg, and early cleavage stage embryo, zinc is the most abundant transition metal compared to iron and copper on average across the 8-cell, morula, and blastocyst (Table 1) (Kim et al., 2010, Kong et al., 2015). While previous studies show increasing copper and zinc content during meiotic maturation in the oocyte, the total number of copper and zinc atoms in the embryo remained constant through late preimplantation embryo development (Table 1 and Fig. 1B) (Kim et al., 2010, Kong et al., 2015). Interestingly, we observed a decrease in total number of iron atoms at the late blastocyst stage compared to 8-cell, morula and early blastocyst (Fig 1B, p<0.05). To illustrate the abundance of zinc atoms in the late preimplantation embryo, we compared the ratio of zinc atom number to copper and iron. The ratio of Zn to Cu is relatively constant and between 16-23 fold higher across all stages analyzed (Table 1). When comparing the ratio of total number of zinc atoms to number of iron atoms, we observed approximately a 2-fold difference in zinc atom abundance at the 8-cell, morula and early blastocyst which doubled to over 4-fold difference in the late blastocyst (Table 1, Fig. 1C, p<0.05).

**Figure 1.**
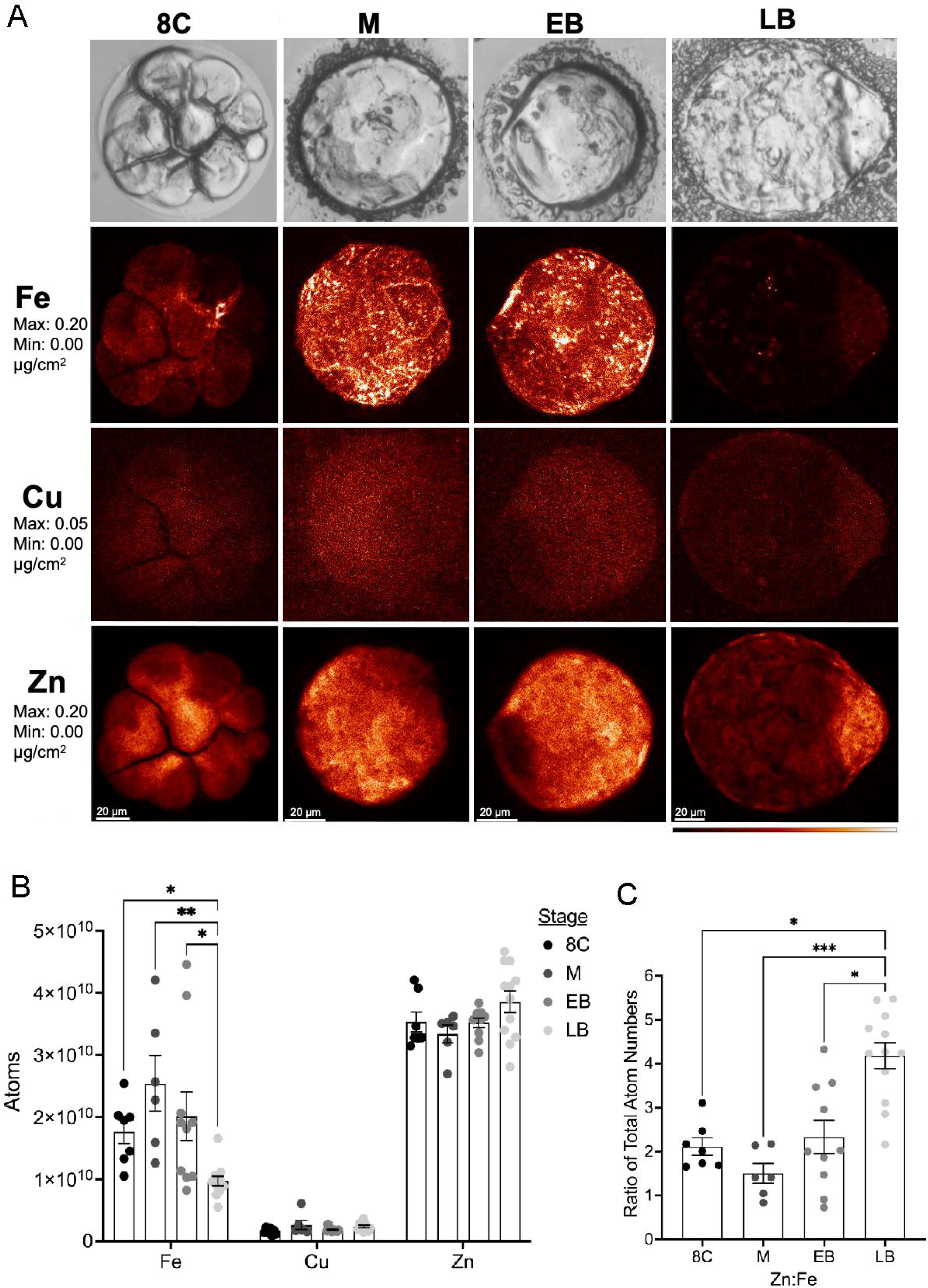
Synchrotron-based X-ray fluorescence microscopy illustrates elemental content of iron, copper and zinc through late preimplantation embryo development. A) Representative whole-mount eight-cell (8C, n=7), morula (M, n=6), early blastocyst (EB, n=10), and late blastocyst (LB, n=12) stage embryos used for XFM. Brightfield images after air-drying and corresponding elemental maps for iron (Fe), copper (Cu) and zinc (Zn) for each stage. Maps for Fe and Zn are shown at 0.00-0.20 and Cu is shown at 0.00-0.05 μg/cm^2^ range. White areas on the elemental maps denote higher atom abundance. B) The mean ± SEM of total number of atoms per embryo for Fe, Cu and Zn. Kruskal-Wallis ANOVA, * p<0.05, **p<0.01. C) The mean ± SEM ratio of total number of atoms between Zn and Fe across all embryo stages. Kruskal-Wallis ANOVA, *p<0.05, ***p<0.001

**Table 1.** Summary of numerical values for total iron, copper, and zinc in the preimplantation embryo by synchrotron-based-x-ray fluorescence microscopy.

| Stage | Units | Fe | Cu | Zn | Zn:Cu | Zn:Fe |
| --- | --- | --- | --- | --- | --- | --- |
| Eight-cell<br>N=7 | Atoms x 10<sup>9</sup> | **17.6 ± 1.9** | **1.65 ± 0.2** | **35.3 ± 1.6** | 23.0 ± 2.5 | 2.2 ± 0.2 |
|  | Atoms x 10<sup>5</sup>/μm<sup>2</sup> | 29.1 ± 2.6 | *2.74 ± 0.3* | 58.6 ± 2.5 |  |  |
| Morula<br>N=7 | Atoms x 10<sup>9</sup> | **25.4 ± 4.5** | **2.61 ± 0.7** | **33.4 ± 1.4** | 15.7 ± 2.3 | 1.5 ± 0.2 |
|  | Atoms x 10<sup>5</sup>/μm<sup>2</sup> | 53.6 ± 9.8 | *5.60 ± 1.7* | 70.6 ± 6.6 |  |  |
| Early Blastocyst<br>N=9 | Atoms x 10<sup>9</sup> | **20.2 ± 3.9** | **1.85 ± 0.1** | **35.2 ± 0.8** | 19.4 ± 0.7 | 2.3 ± 0.4 |
|  | Atoms x 10<sup>5</sup>/μm<sup>2</sup> | *47.0 ± 10.7* | *4.09 ± 0.4* | 78.2 ± 5.1 |  |  |
| Late Blastocyst<br>N=9 | Atoms x 10<sup>9</sup> | **9.72 ± 0.8** | **2.40 ± 0.2** | **38.6 ± 1.7** | 16.7 ± 0.8 | 4.2 ± 0.3 |
|  | Atoms x 10<sup>5</sup>/μm<sup>2</sup> | 13.8 ± 1.7 | *3.29 ± 0.3* | 53.2 ± 4.2 |  |  |

Values in **bold** represent number of atoms x 10<sup>9</sup> ± SEM. Values in *italics* represent the concentration of atoms x 10<sup>5</sup> ± SEM. Zn:Fe and Zn:Cu are the ratios of total number of atoms of zinc to iron or copper ± SEM. Fe=iron, Cu=copper, Zn=Zinc.

### Zinc is enriched in the ICM of the blastocyst

Because of the striking localization of zinc atoms in the inner cell mass of the late blastocyst (Fig. 1A), we performed a more in-depth analysis of the XFM data based on defined regions of interest (Fig. 2A). We confirmed that total zinc level in the late blastocyst is significantly higher than both iron and copper (Table 1, Fig. 2B, p<0.001). Using the phosphorus maps to define regions of interest that had high cell density in the blastocyst, we estimated the metal content of the TE and ICM. In general, we observed the number of iron, copper and zinc atoms per μm^2^ in the ICM (1.72 × 10^6^, 4.07 × 10^5^, 7.82 × 10^6^) was higher compared to the TE (9.38 × 10^5^, 2.97 × 10^5^, 3.86 × 10^6^ atoms/μm^2^) (Fig. 2C, p<0.001). The ratio of ICM to TE metal atoms per area was significantly increased in iron (1.9) and zinc (2.1) compared to copper (1.4) (Fig 2D, p<0.05). Taken together, we observed that zinc is the most abundant transition metal in the blastocyst compared to iron and copper with more than double the concentration of zinc atoms in the ICM.

**Figure 2.**
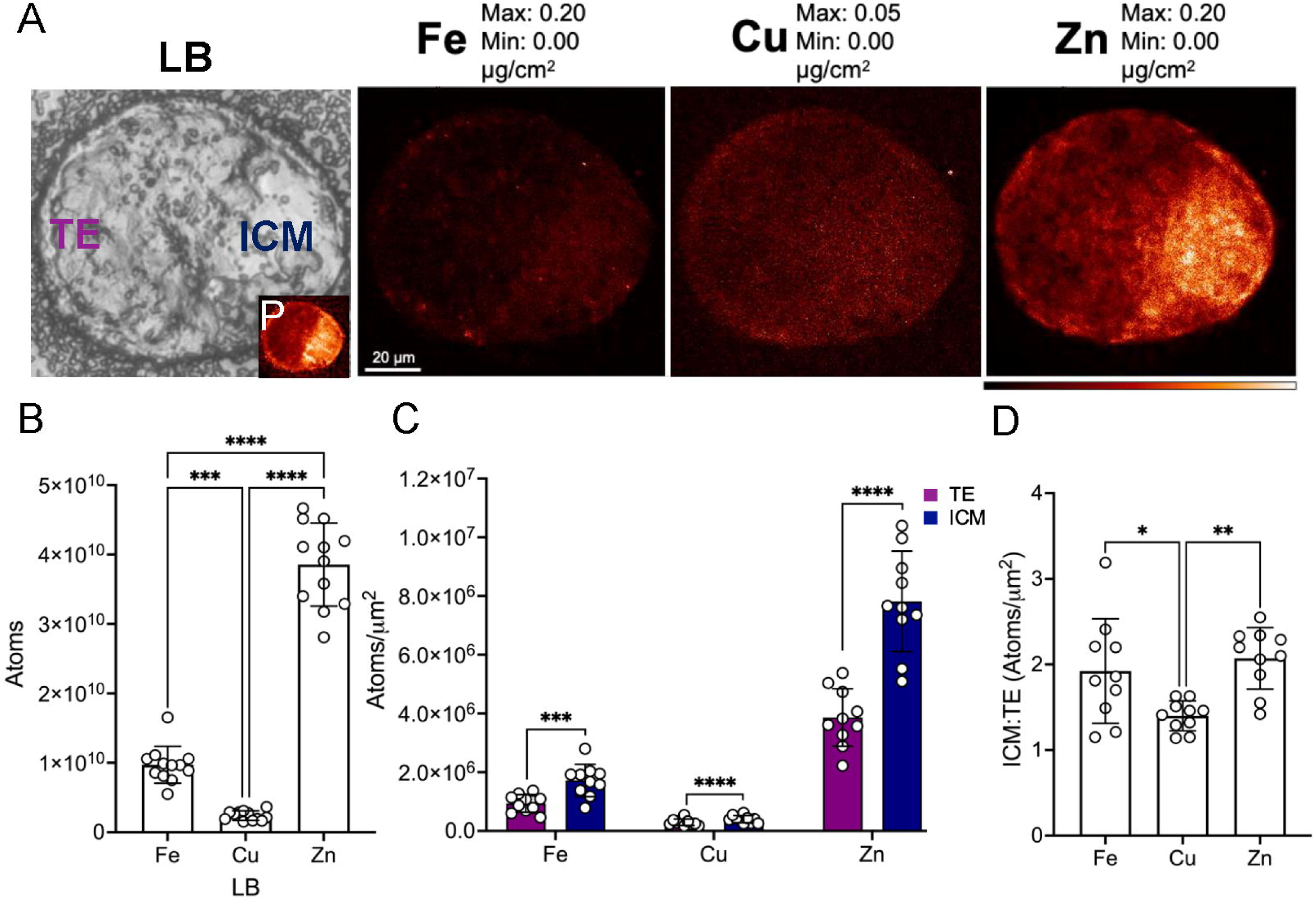
XFM analysis of atoms in the trophectoderm (TE) and inner cell mass (ICM) regions of the late blastocyst. A) Representative brightfield and elemental maps for Fe, Cu and Zn in the late blastocyst. Inset shows phosphorus (P) map (0.00-25.0 μg/cm^2^) which was used to define regions by cell density. Maps for Fe and Zn are shown at 0.00-0.20 and Cu is shown at 0.00-0.05 μg/cm^2^ range. B) Total number of atoms ± SD of Fe, Cu and Zn in the late blastocyst (n=12). Ordinary one-way ANOVA *** p<0.001, ****p<0.0001. C) The number of atoms per μm^2^ ± SD of Fe, Cu and Zn in the TE (purple) and ICM (blue) (n=12 late blastocysts). Paired t-test, *** p<0.001, ****p<0.0001. D) The ratio of atom number per μm^2^ ± SD in the ICM to the TE for Fe, Cu and Zn (n=12 late blastocysts). Ordinary one-way ANOVA, *p<0.05, **p<0.01.

### Zinc homeostasis is required for the morula-to-blastocyst transition

Given that zinc is the most abundant transition metal in the late blastocyst and displays an accumulation in the ICM, which is the site of pluripotent stem cells, we sought to determine the functional significance of zinc fluctuations during the late stages of embryo development. Zinc availability can be limited in biological samples through treatment with intracellular chelators or increased by treatment with zinc ionophores. We have previously shown that zinc availability is essential for mitosis in the cleavage stage embryo, since chelation using TPEN blocks cell cycle progression and precludes development (Kong et al., 2015). Therefore, to determine the role of zinc during the morula-to-blastocyst transition, we perturbed zinc homeostasis in the murine embryo using a model of zinc excess where ZPT was used to elevate intracellular zinc (Fig. 3A) (Taki et al., 2004, Kim et al., 2011, Bernhardt et al., 2012). We first performed a dose-response assay to determine the maximal dose of ZPT that increased zinc without impacting the morula-to-blastocyst transition. To do this, morulae were cultured in control (0 nM), 10 nM, 100 nM and 1 μM of ZPT for 24 hours, and the number of blastocysts or degenerating embryos was scored (Fig. 3B). There was no effect on blastocyst development with 10 nM (85.0 ± 10.0%) or 100 nM (87.1 ± 9.5%) ZPT compared to control (87.5 ± 7.1%) (p>0.05). However, treatment with 1 μM ZPT led to the development of significantly fewer blastocysts (16.0 ± 18.2%) compared to the other doses (Fig. 3C, p<0.05). These results demonstrate embryos readily tolerate as much as 100 nM ZPT during the morula-to-blastocyst transition with no alteration in blastocyst development or change in blastocyst diameter and area compared to control (Supp. Fig. 5A, B). However, ZPT treatment of 1 μM is toxic and impairs blastocyst development.

**Figure 3.**
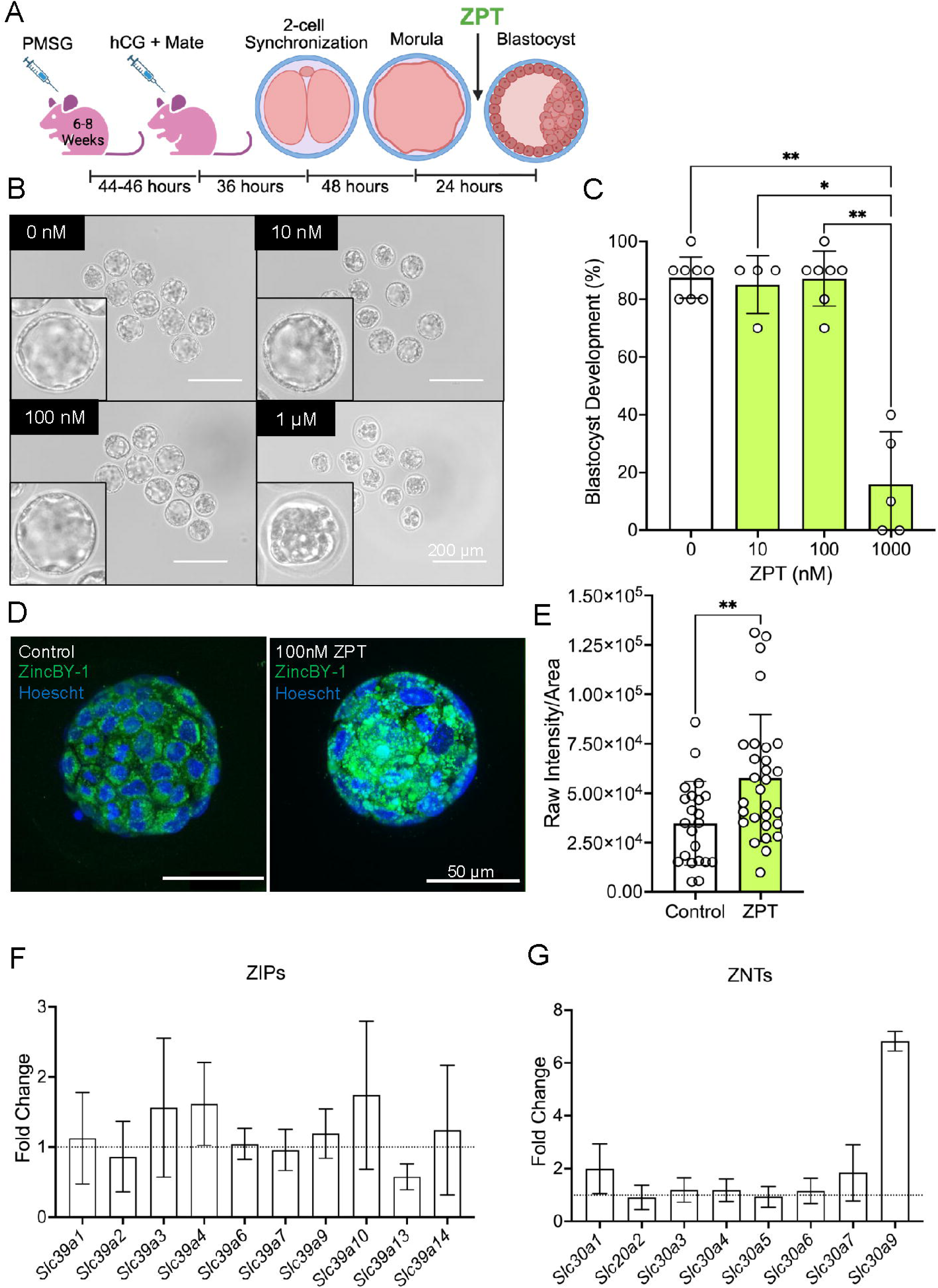
Zinc ionophore, zinc pyrithione (ZPT), treatment during the morula-to-blastocyst transition. A) Experimental paradigm illustrating hyperstimulation and superovulation in CD1 female mice at 6-8 weeks old. Mice were mated and 2-cell embryos were collected and cultured *in vitro* until the morula stage. ZPT treatment was performed at the morula stage and after 24 hours blastocysts were collected for endpoint analysis. Schematic made in Biorender. B) Representative brightfield images of blastocysts cultured after 24 hours in control (0 nM), 10 nM, 100 nM and 1 μM ZPT. Insets display images of representative embryos. B) The percent (mean ± SD) of blastocysts developed from morulae cultured in ZPT for 24 hours. Each datapoint represents the average from a single trial of 10 embryos each. Between 4-8 trials were performed per ZPT dose. Ordinary one-way ANOVA, **p<0.01, *p<0.05. C) Representative images of live blastocysts after ZincBY-1 staining. ZincBY-1 (green) stains labile zinc and Hoechst (blue) stains nuclei. Images are maximum projections of the z-stack. D) ZincBY-1 signal intensity (mean ± SD) of control (n=22) or ZPT treated (n=29) blastocysts. Mann-Whitney test, **p<0.01. E) Gene expression of zinc importers (ZIPs), the *Slc39a* family. Data is expressed as fold change of the ΔΔCT (mean ± SD) in expression compared to control. The average of 3 internal replicates from one trial of 50 blastocysts per control and treated is shown. Data normalized to β*-actin* expression. F) Gene expression of zinc exporters (ZNTs), the *Slc30a* family. Data is expressed as fold change of the ΔΔCT (mean ± SD) in expression compared to control. The average of 3 internal replicates from one trial of 50 blastocysts per control and treated is shown. Data normalized to β*-actin* expression. *Slc39a5, Slc39a8, Slc39a11, Slc39a12, Slc30a8,* and *Slc30a10* were included in the experiment but had no expression and are not shown.

**Figure 4.**
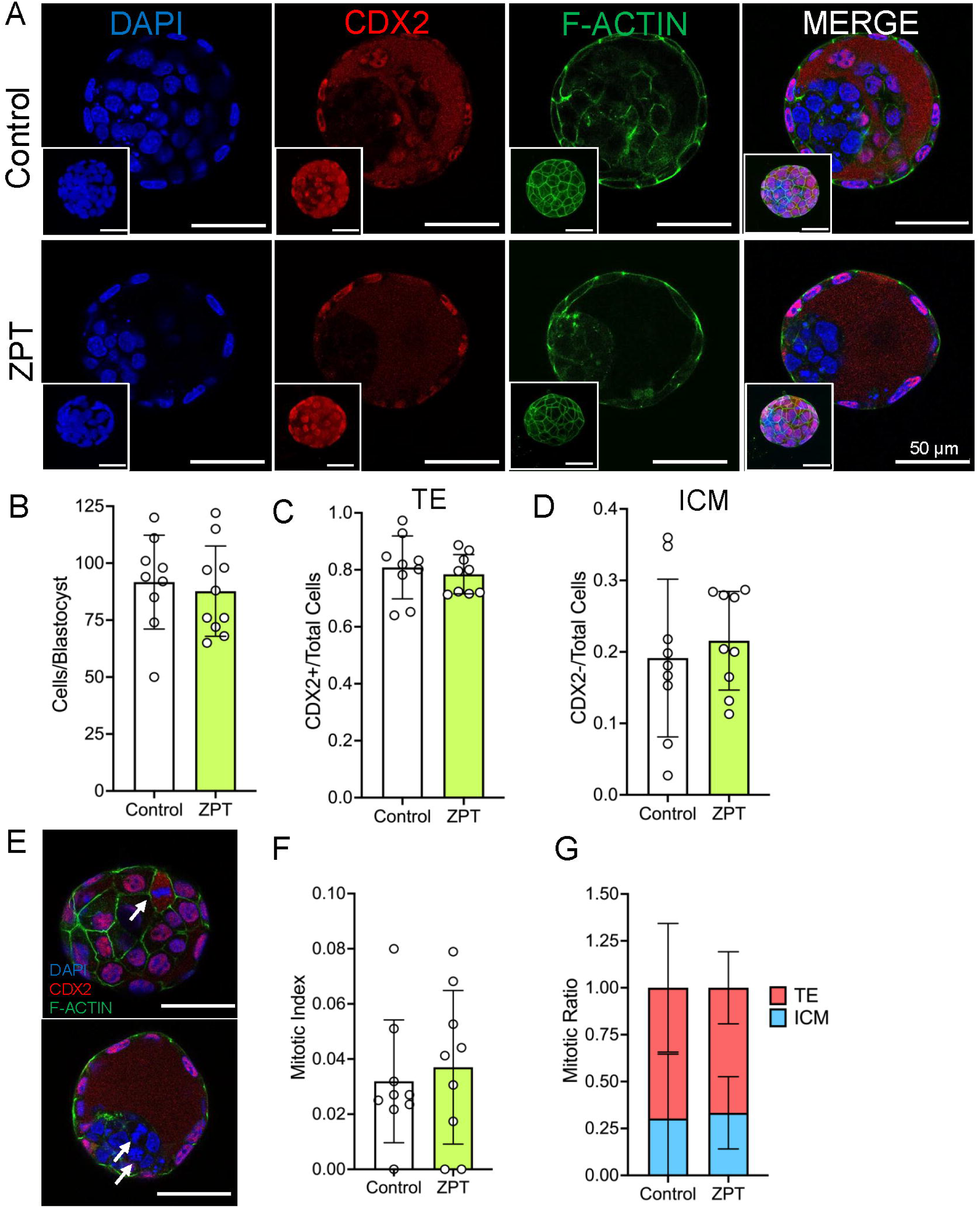
Mitotic endpoints in blastocysts treated with ZPT. A) Representative whole-mount blastocysts of control or ZPT treated blastocysts after immunocytochemistry for CDX2. Mid-sections of blastocysts stained nuclei with DAPI (blue), anti-CDX2 (red), and rhodamine phalloidin (green) and 3 channels merged. Insets show max projection of the z stack. Embryos were generated from a cohort of 10 mice and randomly distributed into treatments groups and each independent assay was repeated at least 2 times. B) The total number of cells per embryo (mean ± SD) were determined by counting nuclei in control (n=9) and treated blastocysts (n=10). Mann-Whitney test, p>0.05. C) The number of CDX2+ nuclei per total nuclei (mean ± SD) determined the TE cells in the embryo (n=9/cohort). Mann-Whitney test, p>0.05. D) The number of CDX2-nuclei per total nuclei (mean ± SD) determined the ICM cells in the embryo (n=9/cohort). Mann-Whitney test, p>0.05. E) Representative images of blastocysts with mitotic cells. Arrows indicate nuclei in mitosis. F) The mitotic index (mean ± SD) captures the nuclei in mitosis per total nuclei per embryo (n=9/cohort). Mann-Whitney test, p>0.05. G) The ratio of the number of mitotic cells (mean ± SD) that are CDX2+ (TE) or CDX2-(ICM) in control (n=9) and ZPT (n=7).

**Figure 5.**
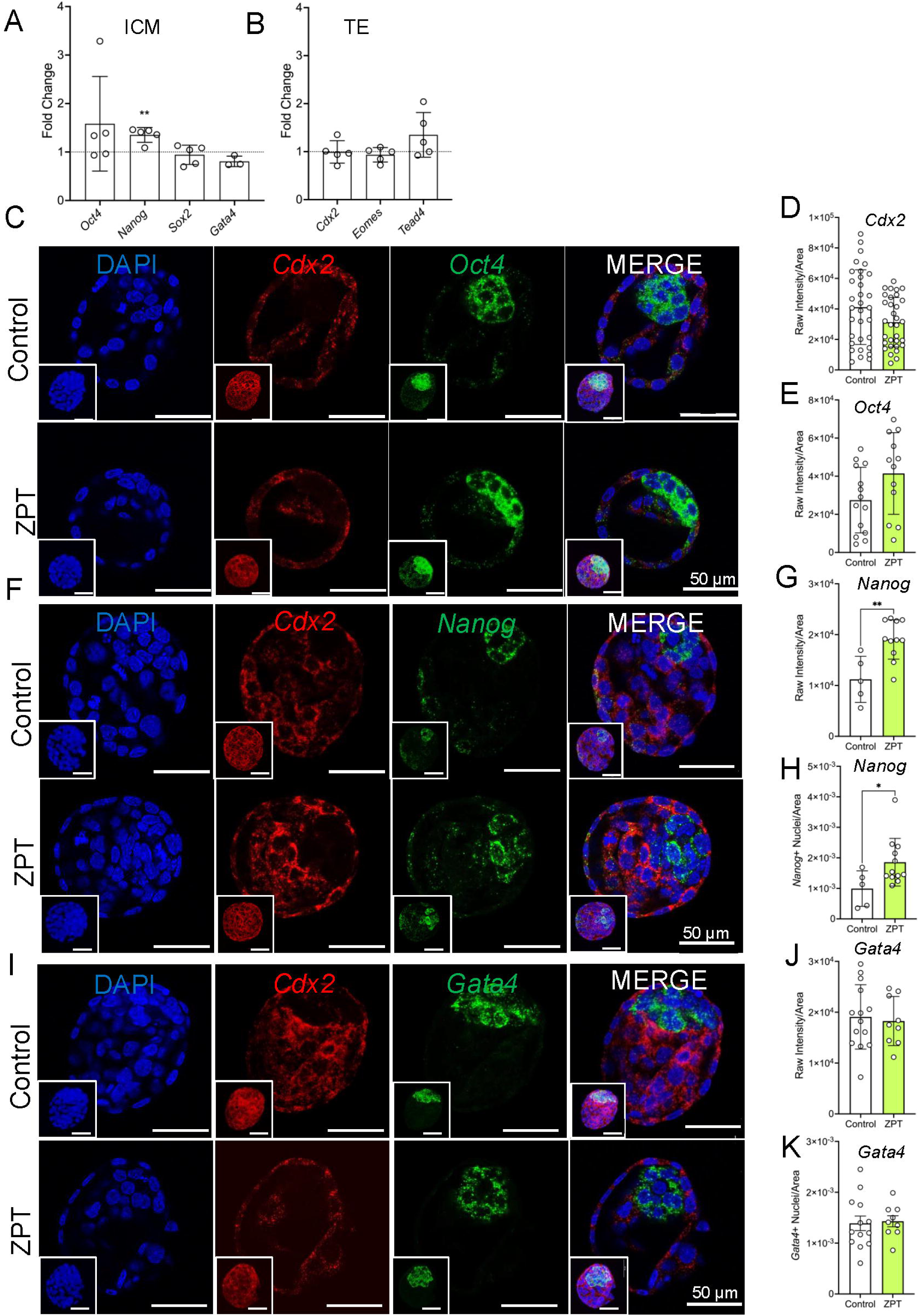
Gene expression of cell fate markers in control and ZPT treated blastocysts. A) Gene expression of inner cell mass genes: *Oct4 (Pouf51), Nanog, Sox2* and *Gata4*. Data is expressed as fold change of the ΔΔCT (mean ± SD) in expression compared to control. Each datapoint represents a trial (n=3-5 trials per gene) of 50 blastocysts per control and treated groups. Three internal replicates were run per trial and averaged. Data normalized to β*-actin* expression. Each gene was assessed by Mann-Whitney test, **p<0.01. B) Gene expression of trophectoderm genes: *Cdx2, Eomes,* and *Tead4*. Data is expressed as fold change of the ΔΔCT (mean ± SD) in expression compared to control. Each datapoint represents a trial (n=5 trials) of 50 blastocysts per control and treated groups. Three internal replicates were run per trial and averaged. Data normalized to β*-actin* expression. Each gene was assessed by Mann-Whitney test, p>0.05. C) Representative mid sections of whole mount blastocysts after fluorescent in situ hybridization for *Cdx2* (red) and *Oct4* (green) transcripts, DAPI (blue) stains nuclei and all channels merged. Each dot represents a single transcript. Inset displays max projection of z stack. D) Transcript intensity per area (mean ± SD) for *Cdx2* per embryo (n=32 embryos/cohort). Mann-Whitney test, p>0.05. Values were pooled from 3 independent trials of randomly sorted embryos derived from 10 mice per trial. E) Transcript intensity per area (mean ± SD) for *Oct4* per embryo (n=14 control and 12 ZPT treated embryos). Mann-Whitney test, p>0.05. Embryos were derived from a cohort of 10 mice and randomly distributed into treatments groups. F) Representative mid sections of whole mount blastocysts after fluorescent in situ hybridization for *Cdx2* (red) and *Nanog* (green) transcripts. G) Transcript intensity per area (mean ± SD) for *Nanog* per embryo (n=5 control and 11 ZPT treated embryos). Mann-Whitney test **p<0.01. Embryos were derived from a cohort of 10 mice and randomly distributed into treatments groups. H) Quantification of nuclei with *Nanog+* transcript per total embryos area in ZPT-treated (n=11) and control (n=5) blastocysts. Mann-Whitney test, *p<0.05. I) Representative mid sections of whole mount blastocysts after fluorescent in situ hybridization for *Cdx2* (red) and *Gata4* (green). J) Transcript intensity per area (mean ± SD) for *Gata4* per embryo (n=14 control and 9 ZPT treated embryos). Mann-Whitney test, p>0.05. Embryos were derived from a cohort of 10 mice and randomly distributed into treatments groups. K) Quantification of *Gata4*+ nuclei per total blastocyst area in control (n=14) and ZPT-treated (n=9) blastocysts. Mann-Whitney test, p<0.05.

### ZPT treatment increases intracellular labile zinc content

While 1 μM of ZPT was cytotoxic to embryos, 100 nM of ZPT did not perturb blastocyst development. To determine whether 100 nM ZPT treatment led to elevation of intracellular zinc levels in treated embryos, we performed live staining with a labile zinc probe, ZincBY-1 (Que et al., 2015). After morulae were cultured for 24 hours in control (0 nM) or 100 nM ZPT and developed into blastocysts, they were then incubated for 30 minutes with ZincBY-1 to visualize labile zinc and Hoechst to visualize nuclei (Fig. 3D). Live blastocysts were imaged with the same exposure settings and the intensity of ZincBY-1 signal was measured. The intensity of ZincBY-1 signal was higher in the ZPT treated blastocysts (5.77 × 10^4^± 3.21 × 10^4^) compared to controls (3.48 × 10^4^ ± 2.21 × 10^4^ raw intensity/area) (Fig. 3E, p<0.05). These findings indicate that ZPT treatment effectively increased intracellular zinc content.

### The zinc transporter network is altered with ZPT treatment

We next assessed whether ZPT treatment during the morula-to-blastocyst transition impacted expression of genes that regulate zinc into and out of cells. We performed RT-qPCR on control or treated blastocysts and analyzed the gene expression of zinc importers (ZIPs), *Slc39a* genes, or zinc exporters (ZNTs), *Slc30a* genes. The ZIP family increases cytosolic zinc by transporting zinc into the cytosol from the extracellular compartment or from intracellular vesicles, whereas the ZNT family reduces cytosolic zinc levels by promoting efflux into the extracellular regions or influx into vesicles (Liuzzi and Cousins, 2004). ZPT treatment at 100nM did not change the expression of the majority of the zinc transport network. However, there was a decreased trend in *Slc39a13 (Zip13*) expression (0.58 ± 0.19) and an increased trend in *Slc30a9 (Znt9)* expression (6.83 ± 0.38 fold change to control) (Fig. 3F and G).

### Excess zinc does not impair mitosis in the blastocyst

Given that zinc is an essential cofactor for mitosis and is necessary to support embryo development, we tested whether increasing intracellular zinc during the morula-to-blastocyst transition impacts the number of resulting blastomeres. Blastocyst cell number is indicative of developmental outcomes and implantation success (Lane and Gardner, 1997, Kong et al., 2016, Guzman et al., 2019). To assess blastocyst cell number, we stained blastocysts with DAPI to count total nuclei during immunocytochemistry (Fig. 4A). There was no difference in total nuclei in control (91.7 ± 20.6) or treated blastocysts (87.7 ± 19.8 nuclei per embryo) (Fig. 4B, p>0.05). Next we tested whether there was a difference in the fraction of TE or ICM cells after ZPT treatment by staining with CDX2, a marker for the TE lineage. There were 0.81 ± 0.11 CDX2 positive nuclei/total nuclei in control embryos and 0.78 ± 0.07 CDX2+ nuclei/total nuclei in ZPT treated embryos (Fig. 4C, p>0.05). CDX2 negative nuclei were counted as a proxy for ICM cells, and there was no difference between control (0.19 ± 0.11) and ZPT treated blastocysts (0.22 ± 0.07 CDX2-nuclei/total nuclei) (Fig. 4D, p>0.05). Thus, elevated zinc during blastocyst development did not impact total numbers of TE or ICM cells. Next, we assessed whether ZPT treatment impacted the mitotic index or the number of cells that were in metaphase or anaphase (Fig. 4E). There was no difference in the mitotic index between control (0.03 ± 0.02) and treated (0.04 ± 0.03 mitotic/total cells) blastocysts (Fig. 4F p>0.05). Lastly to confirm the number of mitotic cells was equivalent in these embryos, we assessed if there was a difference in the ratio of CDX2 positive and negative mitotic cells and found the ratio was the same in control and treated embryos (Fig. 4G, p>0.05). Here, we demonstrate that elevated levels of intracellular zinc do not alter the number of cells in the resulting blastocyst.

### Zinc ionophore treatment during the morula to blastocyst transition alters expression of pluripotency gene, Nanog

Given that the number of TE and ICM cells was similar in the blastocyst with excess zinc, we next tested whether zinc elevation altered expression of key genes involved in maintaining pluripotency in the ICM and TE lineage using RT-qPCR. There was no difference in the fold change expression of *Oct4* (1.58 ± 0.98), *Sox2* (0.94 ± 0.20), and *Gata4* (0.81 ± 0.11) in ZPT treated embryos relative to control embryos (Fig. 5A, p>0.05). However, we did observe an increase in *Nanog* (1.35 ± 0.15) expression compared to control in the blastocyst (Fig. 5A, p<0.05). Genes involved in the TE lineage were unaffected by zinc, as there was no significant fold change in expression for *Cdx2* (0.99 ± 0.23), *Eomes* (0.94 ± 0.15), or *Tead4* (1.3 ± 0.47-fold change) compared to control (Fig. 5B, p>0.05). While RT-qPCR was performed on pooled control and treated blastocysts, we performed orthogonal validation using RNAscope of cell fate genes in whole-mount blastocysts (Fig. 5C-K). This method of fluorescent in situ hybridization allows for visualization of single RNA transcripts, and we used fluorescent intensity to quantify transcript abundance. Consistent with our qPCR data, we observed no difference in *Cdx2* transcript (4.12 × 10^4^ ± 2.45 × 10^4^ and 3.13 × 10^4^ ± 1.62 × 10^4^) or *Oct4* transcript (2.75 × 10^4^ ± 1.72 × 10^4^ and 4.14 × 10^4^ ± 2.14 × 10^4^ raw intensity of transcript/area) abundance between control and treated embryos (Fig. 5C-E, G, p>0.05). There was an increase in *Nanog* transcript abundance with ZPT treatment (1.91 × 10^4^ ± 3.88 × 10^3^) compared to control (1.12 × 10^4^ ± 4.54 × 10^3^ raw intensity/area) (Fig. 5F, G, p<0.05). In addition, there were more *Nanog*+ nuclei per embryo after ZPT treatment (1.86 × 10^−3^ ± 7.82 × 10^−4^) compared to control (9.93 × 10^−4^ ± 5.81 × 10^−4^ *Nanog*+ nuclei/area) (Fig. 5H. p<0.05). There was no difference in the abundance of *Gata4* transcripts between control (1.91 × 10^4^ ± 6.33 × 10^4^) and treated blastocysts (1.83 × 10^4^ ± 4.82 × 10^4^ raw intensity/area) or number of *Gata4*+ nuclei per embryo between treated (1.43 × 10^−3^ ± 3.24 × 10^−4^ *Gata4*+) and control (1.39 ×10^−3^ ± 4.68×10^−4^ *Gata4+* nuclei/area) (Fig. 5-K. p>0.05). Taken together we observed that elevated zinc during the morula-to-blastocyst transition alters the expression, of pluripotency marker, *Nanog,* in blastocyst.

## Discussion

We began this study with an unbiased profiling of elemental content in the late murine preimplantation embryo. By using x-ray synchrotron microscopy to capture the spatial and quantitative distribution of elements in the murine preimplantation embryo, we observed an abundance of zinc content in the inner cell mass of the blastocyst compared to other metals. Specifically, the number of zinc atoms are more than 16 times greater compared to copper and 4 times greater than iron. These novel descriptions of metal content in the blastocyst led us to ask the question if zinc content could impact cell fate determination. To determine whether zinc impact blastocyst development, we utilized a model of excess zinc treatment in mouse embryos and found ZPT treatment increased intracellular zinc without compromising progression to the blastocyst stage. Although cell number was not impacted, zinc treatment enhanced *Nanog* expression and transcript abundance in the blastocyst. Further work is necessary to reveal the role between zinc and regulation of *Nanog* in the preimplantation embryo; however, these results highlight zinc as a potential cofactor for pluripotency regulation in the developing embryo.

Small molecules that perturb zinc availability are key tools in our arsenal for evaluation of zinc roles during development. The chelating agent, TPEN, has been useful in establishing that elevated levels of zinc in the oocyte are required to maintain cell cycle arrest, prohibit meiotic resumption and control cytoplasmic division during polar body extrusion (Kim et al., 2010, Bernhardt et al., 2012). Because zinc is a critical cofactor for mitosis, similar chelation studies cannot be used as easily to investigate zinc homeostasis in the late-stage preimplantation embryo (Kong et al., 2015). While oocytes can withstand 10 μM of the zinc ionophore ZPT with no cytotoxicity, we saw significant reduction in development when morulae were treated with just 1 μM ZPT (Kim et al., 2011). The embryo appears more sensitive to perturbations that elevate zinc availability than oocytes. This difference may arise from the known transcriptional quiescence in the fully grown oocyte compared to the later stage preimplantation embryo.

Zinc is a key regulatory element and signaling molecule in early development, particularly meiotic progression (Bernhardt 2012). The zinc transporters, ZIPs and ZNTs, modulate zinc level and support zinc signaling in the cell by mediating import into and export out of the cell. However, zinc transporters are also present on organelles and vesicles which could have functional impact as zinc is a signaling molecule and the functional domain in Zn-finger, ring-finger and LIM domain proteins (Cassandri et al., 2017). While additional studies are needed to assess transporter persistence and assembly after zinc supplementation, we observed alteration in expression of zinc transport genes, *Zip13* and *Znt9*, in the blastocyst after ZPT treatment (Fig. 3F, G). *Znt9* facilitates zinc export from the mitochondria, a major storage organelle for zinc (Deng et al., 2021). Loss of *Znt9* leads to mitochondrial zinc dyshomeostasis by impacting morphology, membrane potential and ETC activity. Global knockout of *Znt9* leads to embryonic lethality prior to E10.5 in the mouse (Ge et al., 2024). Therefore, we believe the increased trend in *Znt9* expression in the blastocyst after ZPT treatment is likely compensatory to avoid mitochondrial zinc overload. Alternatively, knock out of *Zip13* is not embryonic lethal during mouse development but shows significant abnormality in hard and soft connective tissue development. *Zip13* loss does not impact cellular zinc concentration but displays zinc trapping in vesicles or overload in the Golgi apparatus (Jeong et al., 2012, Fukada et al., 2008). The proposed mechanistic link between development and *Zip13* is due to *Zip13* regulation of BMP/TGF-β signaling and loss of nuclear SMAD proteins which are co-activate transcription factors involved in connective tissue development (Fukada et al., 2008). In the preimplantation embryo, TGF-β signaling is also essential for activation of *Nanog* and maintenance of pluripotency (Xu et al., 2008). The SMADs are downstream mediators of BMP/TGF-B signaling and specifically, the R-SMADs and SMAD4, contain a Zn-binding motif (MH1) necessary for DNA binding (Shi et al., 1998) (Chai et al., 2003), which suggest loss or excess zinc could directly impact SMAD protein-DNA binding. Taken together alterations in the zinc transport network observed in the blastocyst after ZPT could be compensatory to shuttle zinc from the cytoplasm or organelles to avoid overload but they may also be mediating availability of zinc in signaling pathways that maintain pluripotency. Future studies in this area are necessary to confirm *Znt9* and *Zip13* are robustly perturbed after zinc supplementation in embryonic cells. However, our observations together with previous studies, support the role of zinc transporters as regulators of cellular zinc content as well as providing signaling molecules which may drive transcription.

Our study illustrates the short-term impacts of zinc supplementation on blastocyst formation. Future directions for this work should assess if maintenance of *Nanog* expression by zinc can persist into later developmental stages, impact implantation or alter outgrowth behavior and cell identity. Murine embryonic stem cells treated with zinc chloride maintained *Nanog* expression when exposed to retinoid acid or formed into embryoid bodies to induce differentiation (Hu et al., 2016). Further, zinc is emerging as an alternate media factor to maintain pluripotency in embryonic stem cell culture. In feeder free conditions with media supplemented with zinc, embryonic stem cell maintained self-renewal with gene expression profiles of *Oct4, Nanog* and *Klf4* like the traditional Leukemia inhibitory factor (LIF) supplemented media (Mnatsakanyan et al., 2019). The mechanism of zinc regulation of embryonic stem cell reprogramming is still under investigation; however, several zinc finger proteins, *Zfp143* and *Zfp281*, are transcription factors that interact with the promoter region of *Nanog* to modulate stem cell self-renewal or differentiation (Chen et al., 2008, Kim et al., 2008, Wang et al., 2008, Fidalgo et al., 2011). Thus, our work and others support zinc as a co-factor for cell fate programming in embryonic stem cells; however, larger studies which include protein endpoints and immunostaining in addition to gene expression, are necessary to definitively reveal if zinc can direct embryonic cell identity.

An unexpected result in our study was the significant drop in iron content observed at the late blastocyst stage (Fig. 1A, B). Like zinc, iron homeostasis during development is necessary for metabolism, oogenesis, embryo development and reproductive outcomes (Sieber et al., 2016, Ng et al., 2019, Fisher et al., 2021). For example, iron excess results in impaired blastocyst formation with increased apoptosis and ferroptosis rates as iron excess induced loss of mitochondrial membrane potential and ATP generation (Chen et al., 2021). Interestingly, an increase in iron level at the eight-cell stage compared to the oocyte, egg and early cleavage stage embryo was previously observed and hypothesized to be due to an initiation of mitochondrial respiration and glucose utilization in the embryo (Houghton and Leese, 2004, Kong et al., 2015). Our results suggest this signature is lost at the late-blastocyst stage. The metabolic needs and energy requirements of the preimplantation embryo are dynamic where oxidative phosphorylation is the main source of energy during the cleavage stages; however, as compaction and blastocyst formation occurs, glycolysis increases. Within the blastocyst itself, the energy needs differ where TE cells utilize oxidative phosphorylation and the ICM use glycolysis (Lee and Rinaudo, 2024). While the focus of our study was on zinc, our XFM data displays novel changes in iron content in the blastocyst and distribution in the ICM and TE which may serve as preliminary data for hypotheses around iron regulation of cell fate and function and warrant future investigation.

In conclusion, our studies show the elemental content of the preimplantation embryo and indicate a potential role of zinc in regulation of pluripotency, specifically *Nanog* expression, in the murine blastocyst. To our knowledge, this work provides novel XFM analysis of iron, copper and zinc content in the morula and blastocyst and quantitative differences in metal concentration in the inner cell mass and trophectoderm. This work is consistent with previous reports of zinc supplementation promotes *Nanog* expression in mouse embryonic cells.

## Acknowledgements

This study used resources of the Advanced Photon Source, a U.S. Department of Energy (DOE) Office of Science User Facility, operated for the DOE Office of Science by the Argonne National Laboratory under contract no. DE-AC02-06CH11357. We thank Northwestern University’s Feinberg School of Medicine and Master of Reproductive Science and Medicine program for support, resources and expertise needed to bring together this team and execute the study. We thank Dr. Olga Antipova at the Argonne National Laboratory for assistance in acquiring and analyzing the XFM data, and Dr. Andrew Crawford for helpful discussions. We also thank Dr. Yuying Chen and Dr. Andrew Nowakowski for their contributions and resources in experimental planning and execution. Schematic (Fig. 3A) made using Biorender.

## Author Contributions

J.L.B. wrote the proposal and performed the XFM at the Advanced Photon Source, designed and performed the mouse and *in vitro* experiments, analyzed the data, generated the figures and wrote the manuscript. T.V.O. was responsible for contributing to XFM data acquisition and analysis, utilization of the ZincBY-1 probe, and drafting of the manuscript. F.E.D. contributed to experimental design, data analysis and drafting of the manuscript. T.K.W. was responsible for conceiving, drafting and revising the manuscript and handling correspondence.

---

*Supplemental Figure 1. Elemental Maps for Fe, Cu and Zn in the 8-cell embryo (n=7)*.

Fe and Zn maps are shown at a minimum of 0.00 and maximum of 0.20 μg/cm^2^ range. Cu maps are shown at 0.00-0.05 μg/cm^2^. Whiter areas on the elemental maps denote higher atom abundance.

*Supplemental Figure 2. Elemental Maps for Fe, Cu and Zn in the morula(n=6)*.

Fe and Zn maps are shown at a minimum of 0.00 and maximum of 0.20 μg/cm^2^ range. Cu maps are shown at 0.00-0.05 μg/cm^2^. Whiter areas on the elemental maps denote higher atom abundance. * denotes artifact that was excluded from analysis.

*Supplemental Figure 3. Elemental Maps for Fe, Cu and Zn in the early blastocyst (n=10)*.

Early blastocysts were characterized by a blastocoel cavity <50% of embryo area. Fe and Zn maps are shown at a minimum of 0.00 and maximum of 0.20 μg/cm^2^ range. Cu maps are shown at 0.00-0.05 μg/cm^2^. Whiter areas on the elemental maps denote higher atom abundance.

*Supplemental Figure 4. Elemental Maps for Fe, Cu and Zn in the late blastocyst (n=12)*.

Late blastocysts were characterized by a blastocoel cavity >50% of embryo area. Fe and Zn maps are shown at a minimum of 0.00 and maximum of 0.20 μg/cm^2^ range. Cu maps are shown at 0.00-0.05 μg/cm^2^. Whiter areas on the elemental maps denote higher atom abundance.

*Supplemental Figure 5.* Blastocyst diameter and area after ZPT treatment. A) The cross-sectional diameter of blastocysts (n=60/group) after treatment with control (0 nM ZPT) or 100 nM ZPT. Mann-Whitney test p>0.05. B) The area (µm2) of blastocysts (n=60/group) that developed after treatment with control (0 nM) and 100 nM of ZPT. Mann-Whitney test, p>0.05.

*Supplemental Table 1. Primers used for RT-qPCR*

## References

Artus J, Chazaud C. A close look at the mammalian blastocyst: epiblast and primitive endoderm formation. Cell Mol Life Sci. 2014;71(17):3327–38.

Bernhardt ML, Kim AM, O’Halloran TV, Woodruff TK. Zinc requirement during meiosis I-meiosis II transition in mouse oocytes is independent of the MOS-MAPK pathway. Biol Reprod. 2011;84(3):526–36.

Bernhardt ML, Kong BY, Kim AM, O’Halloran TV, Woodruff TK. A zinc-dependent mechanism regulates meiotic progression in mammalian oocytes. Biol Reprod. 2012;86(4):114.

Cassandri M, Smirnov A, Novelli F, Pitolli C, Agostini M, Malewicz M, et al. Zinc-finger proteins in health and disease. Cell Death Discov. 2017;3:17071.

Chai J, Wu JW, Yan N, Massague J, Pavletich NP, Shi Y. Features of a Smad3 MH1-DNA complex. Roles of water and zinc in DNA binding. J Biol Chem. 2003;278(22):20327–31.

Chen X, Fang F, Liou Y-C, Ng H-H. Zfp143 Regulates Nanog Through Modulation of Oct4 Binding. Stem Cells. 2008;26(11):2759–67.

Chen X, Zhou Y, Wu D, Shu C, Wu R, Li S, et al. Iron overload compromises preimplantation mouse embryo development. Reproductive Toxicology. 2021;105:156–65.

Chen YY, Chen S, Ok K, Duncan FE, O’Halloran TV, Woodruff TK. Zinc dynamics regulate early ovarian follicle development. J Biol Chem. 2023;299(1):102731.

de Jonge HJ, Fehrmann RS, de Bont ES, Hofstra RM, Gerbens F, Kamps WA, et al. Evidence based selection of housekeeping genes. PLoS One. 2007;2(9):e898.

Deng H, Qiao X, Xie T, Fu W, Li H, Zhao Y, et al. SLC-30A9 is required for Zn(2+) homeostasis, Zn(2+) mobilization, and mitochondrial health. Proc Natl Acad Sci U S A. 2021;118(35).

Duncan FE, Que EL, Zhang N, Feinberg EC, O’Halloran TV, Woodruff TK. The zinc spark is an inorganic signature of human egg activation. Sci Rep. 2016;6:24737.

Eide DJ. Zinc transporters and the cellular trafficking of zinc. Biochim Biophys Acta. 2006;1763(7):711–22.

Fidalgo M, Shekar PC, Ang YS, Fujiwara Y, Orkin SH, Wang J. Zfp281 functions as a transcriptional repressor for pluripotency of mouse embryonic stem cells. Stem Cells. 2011;29(11):1705–16.

Fisher AL, Sangkhae V, Balusikova K, Palaskas NJ, Ganz T, Nemeth E. Iron-dependent apoptosis causes embryotoxicity in inflamed and obese pregnancy. Nat Commun. 2021;12(1):4026.

Fukada T, Civic N, Furuichi T, Shimoda S, Mishima K, Higashiyama H, et al. The zinc transporter SLC39A13/ZIP13 is required for connective tissue development; its involvement in BMP/TGF-beta signaling pathways. PLoS One. 2008;3(11):e3642.

Ge J, Li H, Liang X, Zhou B. SLC30A9: an evolutionarily conserved mitochondrial zinc transporter essential for mammalian early embryonic development. Cell Mol Life Sci. 2024;81(1):357.

Guzman L, Nunez D, Lopez R, Inoue N, Portella J, Vizcarra F, et al. The number of biopsied trophectoderm cells may affect pregnancy outcomes. J Assist Reprod Genet. 2019;36(1):145–51.

Houghton FD, Leese HJ. Metabolism and developmental competence of the preimplantation embryo. Eur J Obstet Gynecol Reprod Biol. 2004;115 Suppl 1:S92–6.

Hu J, Yang Z, Wang J, Yu J, Guo J, Liu S, et al. Zinc Chloride Transiently Maintains Mouse Embryonic Stem Cell Pluripotency by Activating Stat3 Signaling. PLoS One. 2016;11(2):e0148994.

Jedrusik A. Making the first decision: lessons from the mouse. Reprod Med Biol. 2015;14(4):135–50.

Jeong J, Walker JM, Wang F, Park JG, Palmer AE, Giunta C, et al. Promotion of vesicular zinc efflux by ZIP13 and its implications for spondylocheiro dysplastic Ehlers-Danlos syndrome. Proc Natl Acad Sci U S A. 2012;109(51):E3530–8.

Kim AM, Bernhardt ML, Kong BY, Ahn RW, Vogt S, Woodruff TK, et al. Zinc sparks are triggered by fertilization and facilitate cell cycle resumption in mammalian eggs. ACS Chem Biol. 2011;6(7):716–23.

Kim AM, Vogt S, O’Halloran TV, Woodruff TK. Zinc availability regulates exit from meiosis in maturing mammalian oocytes. Nat Chem Biol. 2010;6(9):674–81.

Kim J, Chu J, Shen X, Wang J, Orkin SH. An extended transcriptional network for pluripotency of embryonic stem cells. Cell. 2008;132(6):1049–61.

Kong BY, Bernhardt ML, Kim AM, O’Halloran TV, Woodruff TK. Zinc maintains prophase I arrest in mouse oocytes through regulation of the MOS-MAPK pathway. Biol Reprod. 2012;87(1):11, 1-2.

Kong BY, Duncan FE, Que EL, Xu Y, Vogt S, O’Halloran TV, et al. The inorganic anatomy of the mammalian preimplantation embryo and the requirement of zinc during the first mitotic divisions. Dev Dyn. 2015;244(8):935–47.

Kong X, Yang S, Gong F, Lu C, Zhang S, Lu G, et al. The Relationship between Cell Number, Division Behavior and Developmental Potential of Cleavage Stage Human Embryos: A Time-Lapse Study. PLoS One. 2016;11(4):e0153697.

Lane M, Gardner DK. Differential regulation of mouse embryo development and viability by amino acids. J Reprod Fertil. 1997;109(1):153–64.

Lee SH, Rinaudo PF. Metabolic regulation of preimplantation embryo development in vivo and in vitro: Molecular mechanisms and insights. Biochem Biophys Res Commun. 2024;726:150256.

Liuzzi JP, Cousins RJ. Mammalian zinc transporters. Annu Rev Nutr. 2004;24:151–72.

Maemura M, Taketsuru H, Nakajima Y, Shao R, Kakihara A, Nogami J, et al. Totipotency of mouse zygotes extends to single blastomeres of embryos at the four-cell stage. Sci Rep. 2021;11(1):11167.

Mamo S, Gal AB, Bodo S, Dinnyes A. Quantitative evaluation and selection of reference genes in mouse oocytes and embryos cultured in vivo and in vitro. BMC Dev Biol. 2007;7:14.

Mnatsakanyan H, Sabater ISR, Salmeron-Sanchez M, Rico P. Zinc Maintains Embryonic Stem Cell Pluripotency and Multilineage Differentiation Potential via AKT Activation. Front Cell Dev Biol. 2019;7:180.

Ng SW, Norwitz SG, Norwitz ER. The Impact of Iron Overload and Ferroptosis on Reproductive Disorders in Humans: Implications for Preeclampsia. Int J Mol Sci. 2019;20(13).

Pfaffl MW, Tichopad A, Prgomet C, Neuvians TP. Determination of stable housekeeping genes, differentially regulated target genes and sample integrity: BestKeeper--Excel-based tool using pair-wise correlations. Biotechnol Lett. 2004;26(6):509–15.

Que EL, Bleher R, Duncan FE, Kong BY, Gleber SC, Vogt S, et al. Quantitative mapping of zinc fluxes in the mammalian egg reveals the origin of fertilization-induced zinc sparks. Nat Chem. 2015;7(2):130–9.

Shi Y, Wang YF, Jayaraman L, Yang H, Massague J, Pavletich NP. Crystal structure of a Smad MH1 domain bound to DNA: insights on DNA binding in TGF-beta signaling. Cell. 1998;94(5):585–94.

Sieber MH, Thomsen MB, Spradling AC. Electron Transport Chain Remodeling by GSK3 during Oogenesis Connects Nutrient State to Reproduction. Cell. 2016;164(3):420–32.

Suwinska A, Czolowska R, Ozdzenski W, Tarkowski AK. Blastomeres of the mouse embryo lose totipotency after the fifth cleavage division: expression of Cdx2 and Oct4 and developmental potential of inner and outer blastomeres of 16- and 32-cell embryos. Dev Biol. 2008;322(1):133–44.

Taki M, Wolford JL, O’Halloran TV. Emission ratiometric imaging of intracellular zinc: design of a benzoxazole fluorescent sensor and its application in two-photon microscopy. J Am Chem Soc. 2004;126(3):712–3.

Tian X, Anthony K, Neuberger T, Diaz FJ. Preconception zinc deficiency disrupts postimplantation fetal and placental development in mice. Biol Reprod. 2014;90(4):83.

Uriu-Adams JY, Keen CL. Zinc and reproduction: effects of zinc deficiency on prenatal and early postnatal development. Birth Defects Res B Dev Reprod Toxicol. 2010;89(4):313–25.

Wang ZX, Teh CH, Chan CM, Chu C, Rossbach M, Kunarso G, et al. The transcription factor Zfp281 controls embryonic stem cell pluripotency by direct activation and repression of target genes. Stem Cells. 2008;26(11):2791–9.

Willems E, Mateizel I, Kemp C, Cauffman G, Sermon K, Leyns L. Selection of reference genes in mouse embryos and in differentiating human and mouse ES cells. Int J Dev Biol. 2006;50(7):627–35.

Wu G, Gentile L, Fuchikami T, Sutter J, Psathaki K, Esteves TC, et al. Initiation of trophectoderm lineage specification in mouse embryos is independent of Cdx2. Development. 2010;137(24):4159–69.

Xie F, Timme KA, Wood JR. Using Single Molecule mRNA Fluorescent in Situ Hybridization (RNA-FISH) to Quantify mRNAs in Individual Murine Oocytes and Embryos. Sci Rep. 2018;8(1):7930.

Xu RH, Sampsell-Barron TL, Gu F, Root S, Peck RM, Pan G, et al. NANOG is a direct target of TGFbeta/activin-mediated SMAD signaling in human ESCs. Cell Stem Cell. 2008;3(2):196–206.

Zernicka-Goetz M, Morris SA, Bruce AW. Making a firm decision: multifaceted regulation of cell fate in the early mouse embryo. Nat Rev Genet. 2009;10(7):467–77.

Zigo M, Kerns K, Sen S, Essien C, Oko R, Xu D, et al. Zinc is a master-regulator of sperm function associated with binding, motility, and metabolic modulation during porcine sperm capacitation. Commun Biol. 2022;5(1):538.

