## Supplementary material for "Inorganic profiles of preimplantation embryos and the association of zinc with *Nanog* expression in the blastocyst": Supp Table 1

Supplemental Table 1. Primers for RT-qPCR

| Gene | Primer | Sequence (5'-3') | Gene | Primer | Sequence (5'-3') |
|---|---|---|---|---|---|
| *B-actin* | F | CCTGAGGCTCTTTTCCAGCC | *Slc39a9* | F | GCATTAGAGGCAGCAGGAAC |
| | R | TAGAGGTCTTTACGGATGTCAAC | | R | GCATTAAGGCATCCACACCT |
| *Cdx2* | F | AAACCTGTGCGAGTGGATG | *Slc39a10* | F | TACCCACCAGCTTTTCACA |
| | R | TCTGTGTACACCACCCGGTA | | R | TCACTGTGAGCAACGGAGTC |
| *Oct4* | F | TTGGGCTAGAGAAGGATGTGGTT | *Slc39a11* | F | CTTCTTCACCTGGGCAATGT |
| | R | GGAAAAGGGACTGAGTAGAGTGT | | R | GGAGGTCAGCCAGGTAGACA |
| *Nanog* | F | GGTTGAAGACTAGCAATGGTCTG | *Slc39a12* | F | GACTGCAAGCTGTGTTTGGA |
| | R | TGCAATGGATGCTGGGATACT | | R | CTAAGGCCGAGTAGGCTGTG |
| *Gata4* | F | GGGCTGTCATCTCACTATGGGCA | *Slc39a13* | F | GCCTGTCGCCTGGATAATAA |
| | R | TGATTATGTCCCCATGACTGTCAG | | R | CCACCTAAGGCAAAGCTGAG |
| *Sox2* | F | GCGGAGTGGAAACTTTTGTCC | *Slc39a14* | F | TCAGCCGTGTGCTCACTTAC |
| | R | GGGAAGCGTGTACTTATCCTTCT | | R | GGTGCTCGTTTTTCTGCTTC |
| *Eomes* | F | GTGACAGAGACGGRGTGGAGG | *Slc30a1* | F | GCTCTCGAGTTGGTCCTGTC |
| | R | AGAGGAGGCCCGTTGGTCTGG | | R | GCCTCATGGTGAGGTAGGAA |
| *Tead4* | F | GCACCATTACCTCCAACGAG | *Slc30a2* | F | TTCTGGAAGTCACCCTGACC |
| | R | GATCAGCTCATTCCGACCAT | | R | CTAATGAGCATGCTGGCAAA |
| *Slc39a1* | F | GGTCTCTCTGCCAGTTTTCG | *Slc30a3* | F | CCATCAGCACCTTCCTCTTC |
| | R | CAGCATTAAGGAGGCAGAGG | | R | ATGGAGATCATGGGTTGCTC |
| *Slc39a2* | F | CCTGCTTGCTCTTCTGGTTC | *Slc30a4* | F | TGCCGTCCTCTACTTGCTTT |
| | R | CCTCCAGAGCTTCAGCAGTC | | R | TAGGCGATGAAATCCAAAGG |
| *Slc39a3* | F | CCATGGTTCACACACAGAGG | *Slc30a5* | F | TTGGTTTTCATACGGCTTCC |
| | R | AGGGTCCCTGAGGTCACTTT | | R | TTTGGACACGTCCATTTTGA |
| *Sc39a4* | F | CTTGGCTCTAGGCAAACCTG | *Slc30a6* | F | GTCCACGCTGACTGTTCAGA |
| | R | AGTGTGGCCAGGTAATCGTC | | R | CGTTTTTCCCAGGCGTATTA |
| *Slc39a5* | F | GCCAGAGGGAGAACAGACAG | *Slc30a7* | F | GCCACCATACCGAGTCACTT |
| | R | GTGGCAGAAGACTGCTAGGG | | R | CCACAAAAGCGAAAGAGAGG |
| *Slc39a6* | F | TTCCTGTCTCTGCTGGGAGT | *Slc30a8* | F | ACTGATGCGGCTCATCTCTT |
| | R | TGTGCTGATGACTTGCATGA | | R | GATGCAAAGGACAGACAGCA |
| *Slc39a7* | F | AGGAGTGTCAGCCTTGGAGA | *Slc30a9* | F | GTCACCCACGGTCTCTCATT |
| | R | ATTAGGGACCATCGGGTAGG | | R | ACTGCCTGTTCCAAAACCAC |
| *Slc39a8* | F | AGCCTAACGGACACATCCAC | *Slc30a10* | F | TGTGCATGCTAAGGAACTGC |
| | R | AGTACAAGATGCCCCAATCG | | R | CACGGTCCAAGAATGGACTT |
