## Supplementary material for "Inorganic profiles of preimplantation embryos and the association of zinc with *Nanog* expression in the blastocyst": Supp Fig 1-5

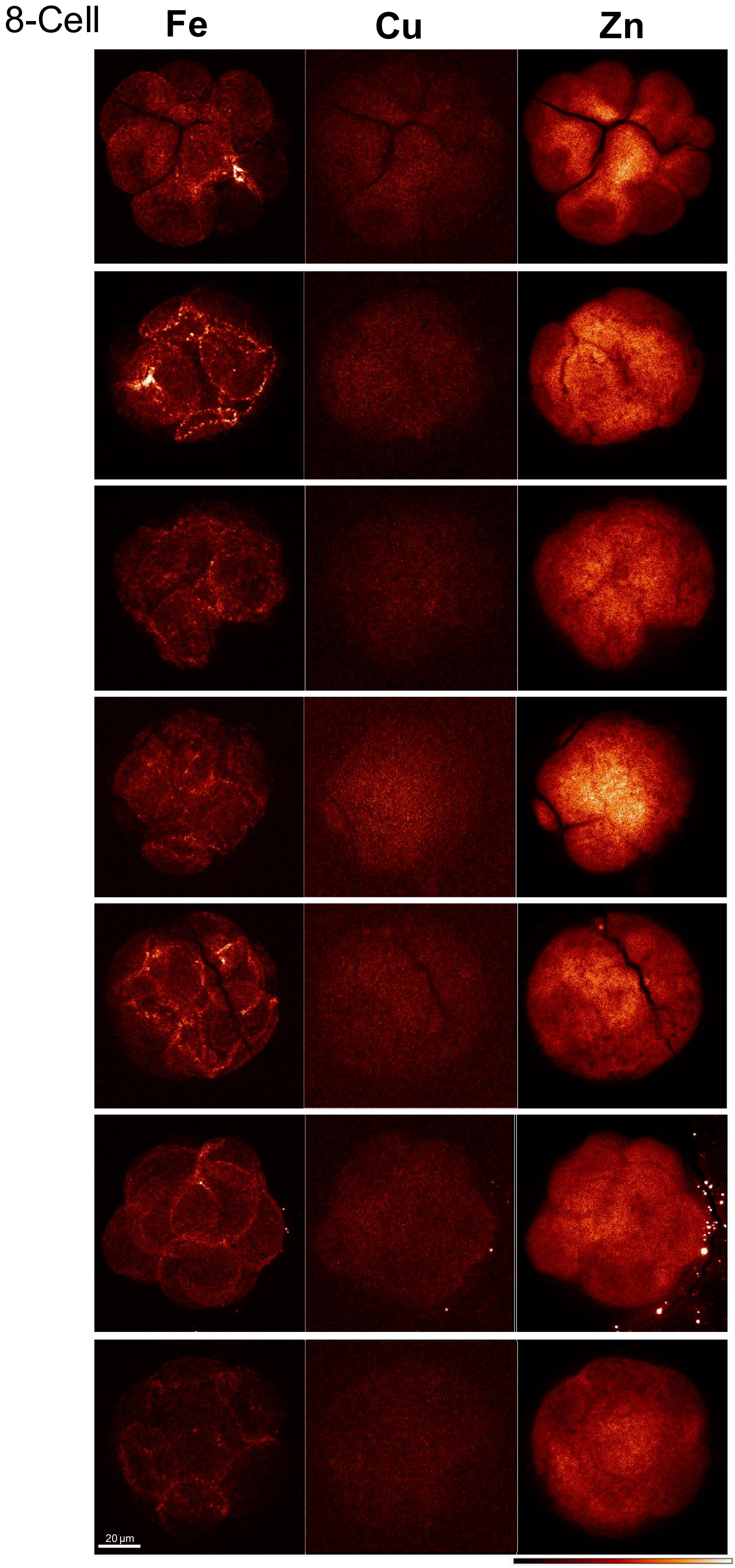


Supp. Fig 1. Elemental Maps for Fe, Cu and Zn in the 8-cell embryo (n=7). Fe and Zn maps are shown at a minimum of 0.00 and maximum of 0.20 µg/cm$^2$ range. Cu maps are shown at 0.00-0.05 µg/cm$^2$. Whiter areas on the elemental maps denote higher atom abundance.

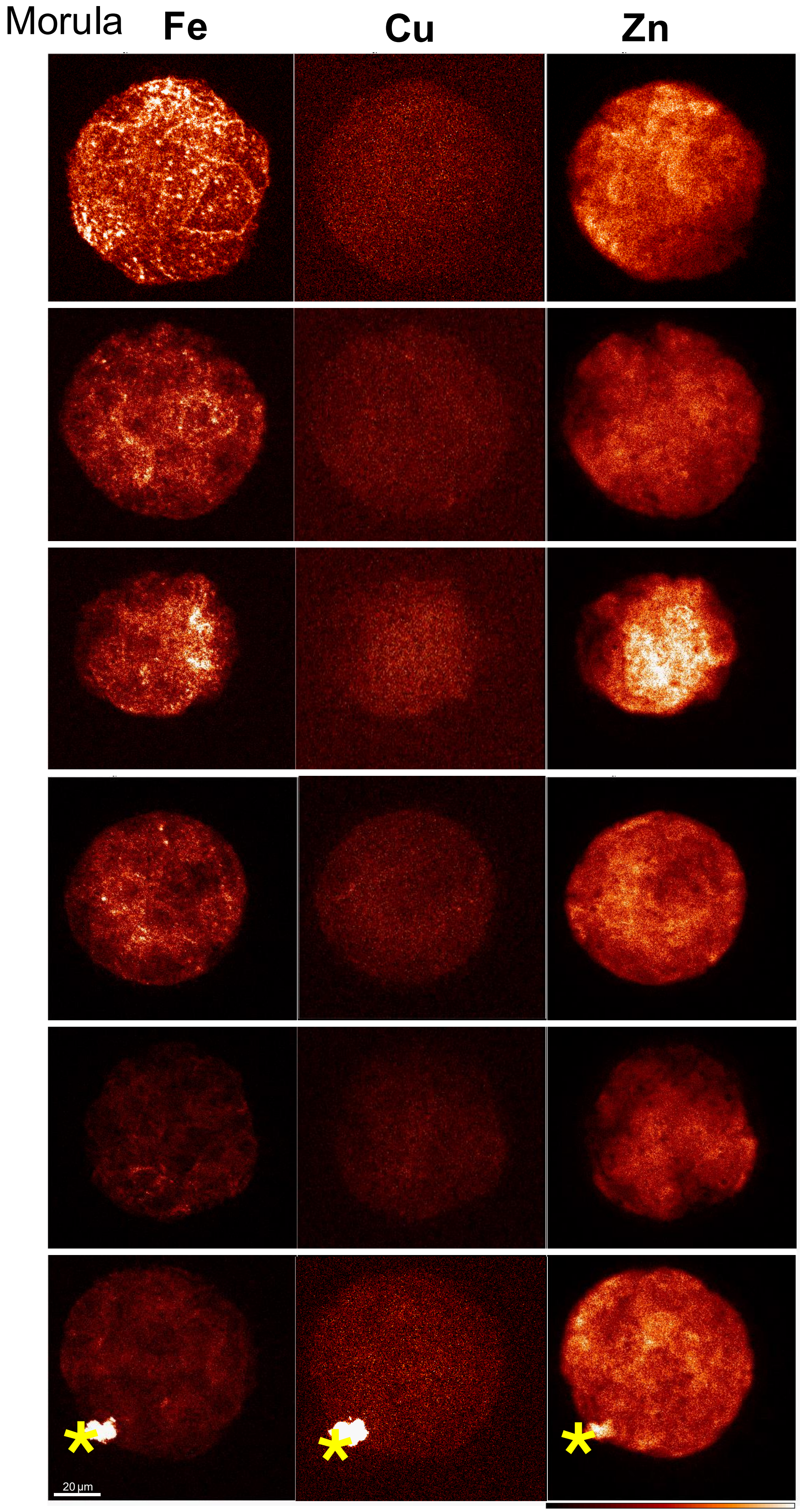


Supp. Fig. 2. Elemental Maps for Fe, Cu and Zn in the morula(n=6). Fe and Zn maps are shown at a minimum of 0.00 and maximum of 0.20 µg/cm$^2$ range. Cu maps are shown at 0.00-0.05 µg/cm$^2$. Whiter areas on the elemental maps denote higher atom abundance. * denotes artifact that was excluded from analysis.

### Early Blastocyst

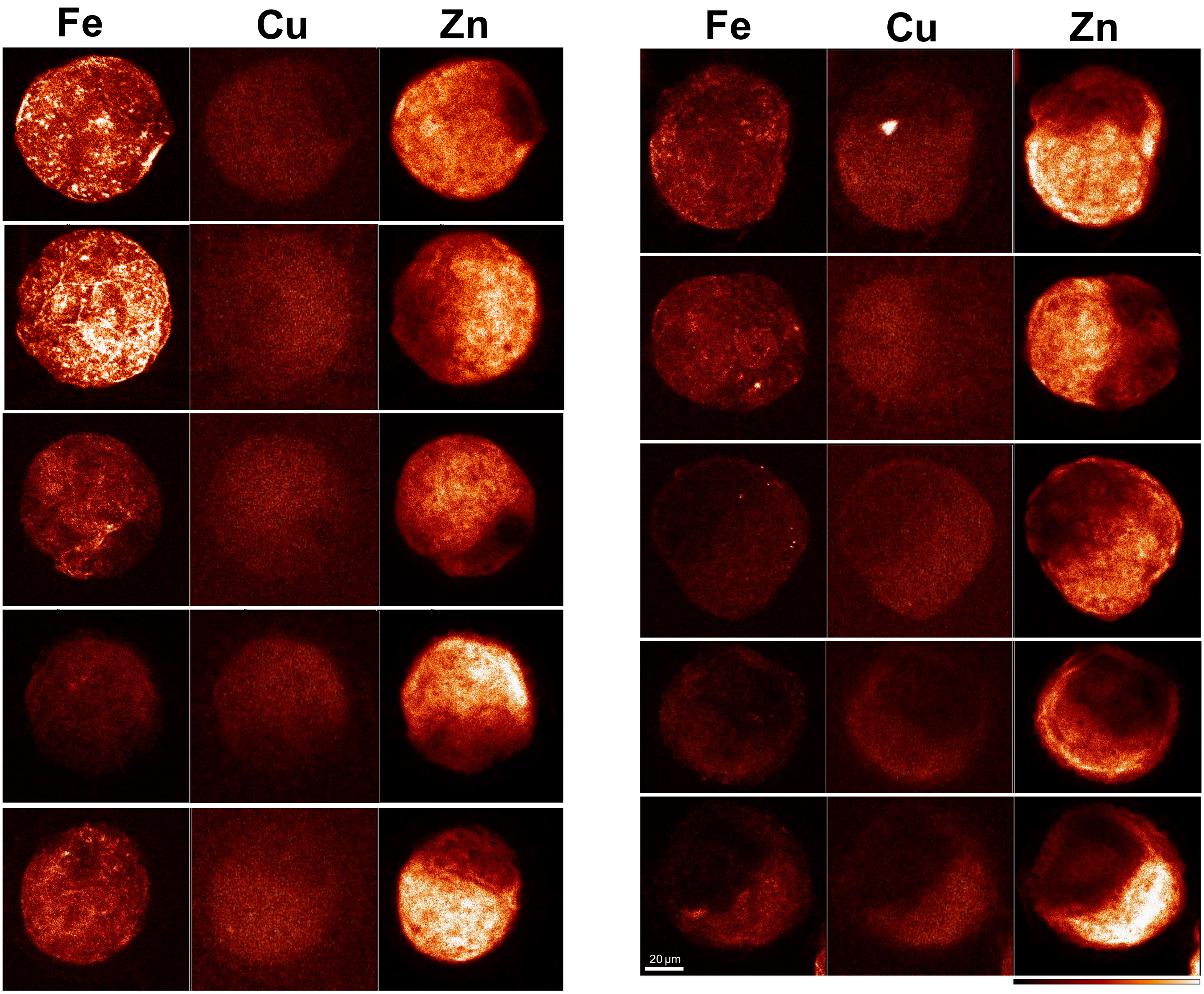


Supp. Fig. 3. Elemental Maps for Fe, Cu and Zn in the early blastocyst (n=10). Early blastocysts were characterized by a blastocoel cavity <50% of embryo area. Fe and Zn maps are shown at a minimum of 0.00 and maximum of 0.20 µg/cm$^2$ range. Cu maps are shown at 0.00-0.05 µg/cm$^2$. Whiter areas on the elemental maps denote higher atom abundance.

### Late Blastocyst

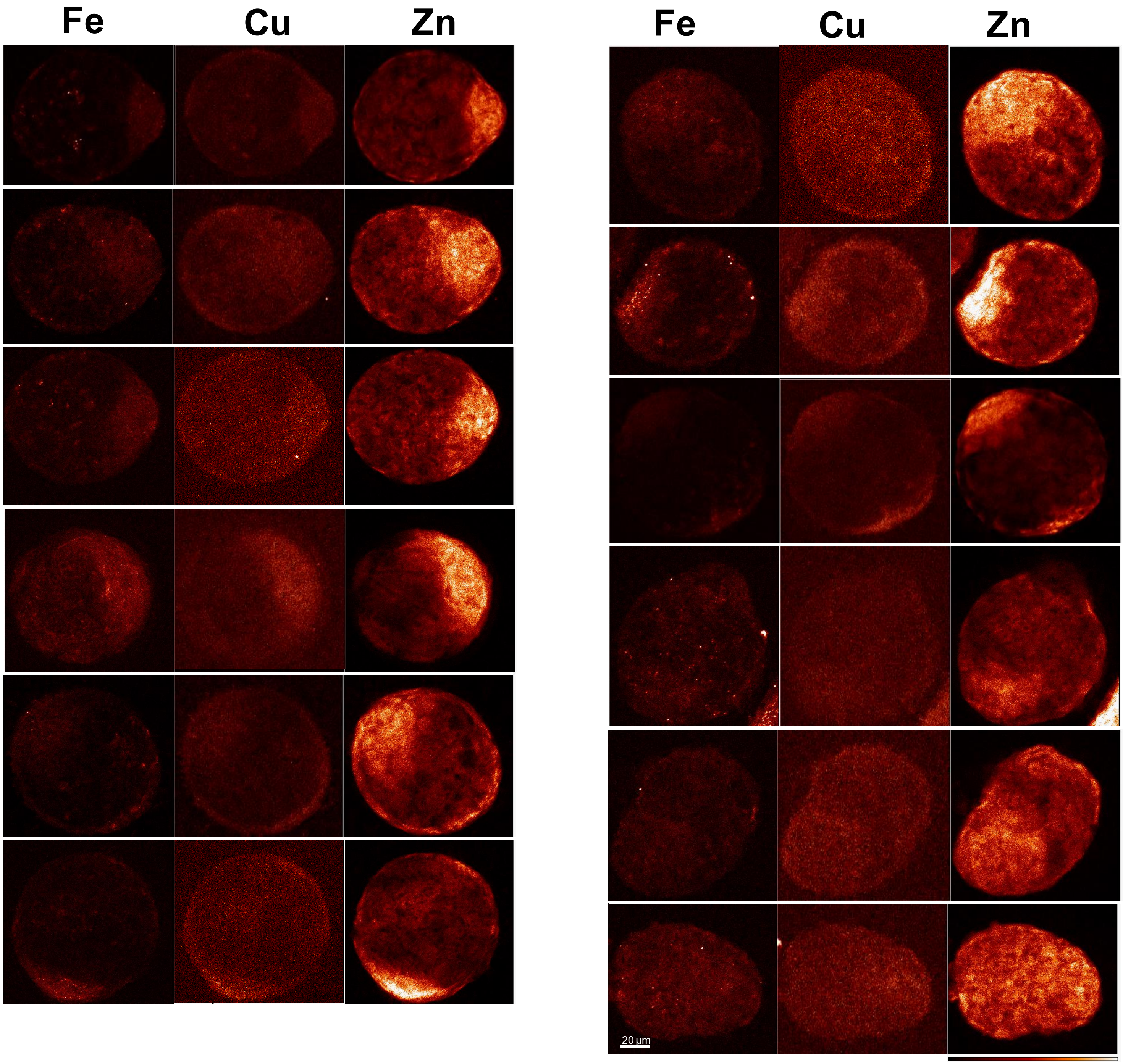


Supp. Fig. 4. Elemental Maps for Fe, Cu and Zn in the late blastocyst (n=12). Late blastocysts were characterized by a blastocoel cavity >50% of embryo area Fe and Zn maps are shown at a minimum of 0.00 and maximum of 0.20 µg/cm$^2$ range. Cu maps are shown at 0.00-0.05 µg/cm$^2$. Whiter areas on the elemental maps denote higher atom abundance.

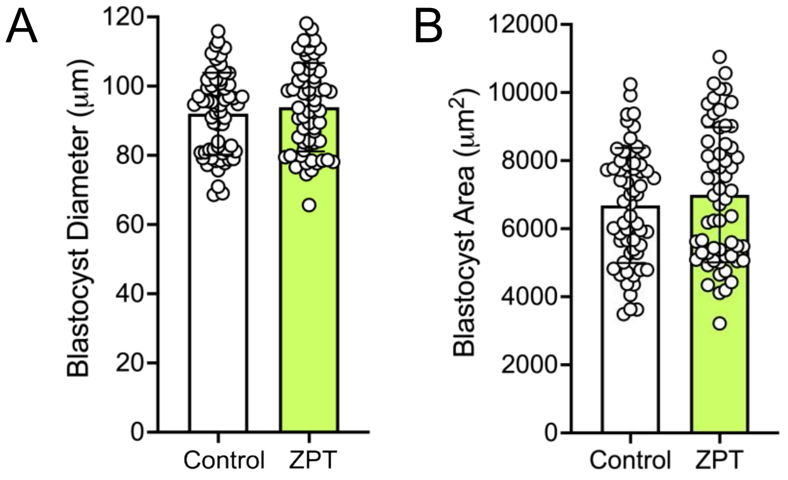


Supp. Fig. 5. Blastocyst diameter and area after ZPT treatment. A) The cross sectional diameter of blastocysts (n=60/group) after treatment with control (0 nM ZPT) or 100 nM ZPT. Mann-Whitney test p>0.05. B) The area (µm2) of blastocysts (n=60/group) that developed after treatment with control (0 nM) and 100 nM of ZPT. Mann-Whitney test, p>0.05.
